# Comparison of place field detection methods and their effect on place field stability and drift in mouse dCA1

**DOI:** 10.64898/2026.03.02.708942

**Authors:** Vladislav Ivantaev, Alireza Chenani, Alessio Attardo, Christian Leibold

## Abstract

**Background:** Hippocampal place cells (PCs) undergo representational drift, i.e., a gradual change in their place fields despite unaltered behavior. While Ca2+ imaging enables long-term tracking of PC populations, distinct PC detection methods have been shown to yield different subpopulations of PCs, with only a few systematic comparisons between methods, especially in open arenas.

**New Method:** We provide an analysis protocol for one-photon PC data obtained during free foraging in two-dimensional arenas that allows us to compare two widely used PC detection methods, significance of spatial information (SI), and split-half correlation (SHC), and their effect on representational drift. The analysis is demonstrated on previously published Ca2+ data from dorsal CA1 of freely foraging mice, with cells tracked for 10 consecutive days.

**Results:** Both criteria, SI and SHC, yielded proportions of approx. 17% PCs with only 40% overlap. SI-identified PCs demonstrated higher stability, higher rate map correlations, and a slower rate of representational drift than SHC-PCs.

**Comparison with existing methods:** Previous studies comparing SI and SHC PC detection methods in Ca2+ data did not focus on either open field behavior or representational drift.

**Conclusion:** Our results indicate that the choice of PC detection method significantly affects the estimate of representational drift in Ca2+ imaging studies.

## 1. Introduction

The hippocampus is critically involved in spatial memory and navigation (O’Keefe et al., 1978), with subfield CA1 directly projecting spatially selective outputs to neocortical areas (Goode et al., 2020). The hippocampal spatial code is prominently carried by CA1 place cells (PCs), which selectively fire when an animal is at a specific location, called the place field (PF) (O’Keefe et al., 1978). Interestingly, the firing patterns of PCs are unstable over time (Mankin et al., 2012; Ziv et al., 2013), despite the stability of the spatial context, a phenomenon referred to as representational drift (Driscoll et al., 2022). Representational drift can occur on a scale of hours to weeks (Ziv et al., 2013; Kinsky et al., 2018), and - besides hippocampus (Mankin et al., 2012; Hainmueller and Bartos, 2018) - it has also been observed in posterior parietal cortex (Driscoll et al., 2017), visual cortex (Deitch et al., 2021), and other sensory brain areas (Aschauer et al., 2022; Schoonover et al., 2021). This is a surprising phenomenon, difficult to reconcile with the ability of the brain to maintain consistent representations over time (Rule et al., 2019). Specifically, in the hippocampus, it is unclear how long-term information storage, generalization, and dynamic memory update coexist with a constant drift of neuronal representations and how this can be beneficial for spatial navigation.

To study representational drift, the same neurons must be tracked over long periods, which can be technically challenging with electrophysiological methods. Optical imaging, by comparison, allows not only to detect neuronal activity- by using Ca^2+^ transients as a proxy in both mice (Ziv et al., 2013; Schoenfeld et al., 2021) and rats (Wirtshafter and Disterhoft, 2022) - but also to track activity patterns over many days up to several weeks (Heiser et al., 2025; Keinath et al., 2022). In fact, optical imaging enables identification of several hundred neurons with micrometer-scale spatial resolution, making it more robust to micrometer-scale tissue motion and drift over time. Thus, recent experimental studies of representational drift rely almost exclusively on calcium imaging as the recording method of choice (Climer et al., 2025; Khatib et al., 2023; Geva et al., 2023).

Crucially, the choice of the behavioral paradigm also plays an important role in studying representational drift and stability of spatial coding. “One-dimensional” linear tracks (Ziv et al., 2013; Wirtshafter and Disterhoft, 2022; Dombeck et al., 2010; Etter et al., 2020; Dong et al., 2021) enable obtaining dense spatial sampling consistent across different sessions, while “two-dimensional” (2D), unconstrained spatial enclosures (Stefanini et al., 2020; Keinath et al., 2022; Chenani et al., 2022; Wang et al., 2024) lead to sparser sampling with reduced consistency across sessions and are thus used less often. In fact, unconstrained (2D) navigation is significantly less biased towards linear sequences of active neurons, thereby drastically reducing spatial averaging within a single session and leading to a dramatic decrease in consistency across sessions. Therefore, studying representational drift in 2D behaviors by using Ca^2+^ imaging requires more sensitive and advanced methods to analyze the spatial selectivity of neurons with complex activity patterns.

One method generally applied to study place coding of neurons is based on the mutual information between position and activity (Ziv et al., 2013). This spatial information (SI), however, was originally derived under the assumption of Poisson spiking (Skaggs et al., 1992), and it is unclear how well it generalizes to Ca^2+^ imaging data. Alternative methods for PC detection (Dombeck et al., 2010; Tanni et al., 2022) have been proposed, most notably a method that identifies PCs *via* the significance of correlation of rate maps derived from two halves of a session (split-half correlation; SHC) (Keinath et al., 2022). However, only a few systematic comparisons of the effects of PC detection methods on representational drift estimates exist (Grijseels et al., 2021), particularly for 2D spatial behaviors.

In this work, we studied the effects of two PC detection methods on the basic properties of CA1 hippocampal PCs during free repeated exploration of a 2D environment using a previously published data set (Chenani et al., 2022). Specifically, we compared the SI- and SHC-identified PCs in terms of the temporal stability of their neural codes. Both PC detection methods provided a similar, but only partially overlapping, set of PCs; however, the SI method yielded more stable PCs than the SHC method. The results further indicate the importance of the PC detection method in studies of representational drift.

## 2. Methods

### 2.1 Data acquisition

The data analyzed here have been published previously (Chenani et al., 2022). The original publication provides a detailed description of the methods. In brief, a group of male mice between three and six months of age was injected with 500 nL of GCaMP6 viral suspension. Later, the mice were implanted with a metal cannula fixed and sealed to their skulls (Attardo et al., 2018).

Before imaging, subjects were allowed to familiarize themselves with the circular arena (47 cm in diameter) for three days, five to ten minutes per day. Then, for 14 days, mice performed free exploration of the arena for at least 15 minutes, while Ca^2+^ transients were recorded using commercial software (Doric Neuroscience Studio) and a continuous wavelength laser (Thorlabs) with the help of a miniature microscope (Doriclenses, S model) at a sampling rate of 45 Hz. To motivate exploration, mixed oat flakes were placed in the bedding material.

### 2.2 Data preprocessing

The exploration behavior of a subject was quantified by binning the trajectory data into a three-dimensional state grid, corresponding to locations on a square grid of 1.8 cm and running direction binned with 90° steps. Sessions were considered for further analysis if more than 30% of all the 3D state bins were occupied for more than 0.5s.

Raw movies were spatially down-sampled by a factor of two and later motion-corrected by the NoRMCorre algorithm (Giovannucci et al., 2019). With the help of the extended constrained non-negative matrix factorization (CNMF-E) algorithm (Pnevmatikakis and Giovannucci, 2017), the Ca^2+^ signal was denoised, deconvolved, and demixed. The method utilizes sparse non-negative deconvolution (Vogelstein et al., 2010) to estimate the starting point of the GCaMP6f fluorescence. The parameters used in the CMNF algorithm are reported in **Supplementary Table 1**. Before being applied to the full session recording, the algorithm was used for the first four consecutive minutes. If the algorithm was able to detect at least 50 ROIs on this reduced recording, the algorithm was applied to the full session. The summary of all sessions discarded and considered across all the subjects used in this work is provided in **Supplementary Table 2**. Only traces with signal-to-noise ratios above 4 and footprints with spatial correlation above 0.05 were subject to place field analysis. Additional individual cells were discarded in case visual inspection revealed saturated spatial pixels or a strongly varying transient baseline.

The algorithm extracts Ca2+ of two kinds: denoised calcium transients and deconvolved spike-like activity. In our work, unless specified otherwise, we have used the denoised calcium transients.

### 2.3 Place field analysis

For all running trajectories, the calcium event locations have been linearly interpolated and smoothed with a 2D Gaussian kernel with σ=4.5 cm. Only the activity samples occurring at running velocities of more than 5 cm/s have been considered in the place field calculation. The cumulative mean activity for each spatial bin x_j_ (place field) pf(x_j_) was determined as follows:

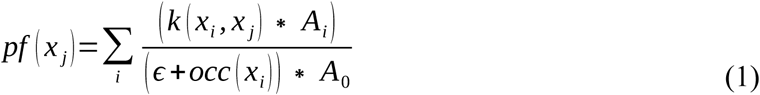

where A_i_ is an amplitude of a DF/F trace at a temporal sample i, 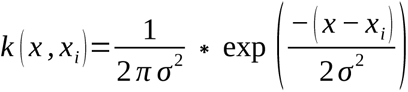 is a normalized Gaussian centered at an interpolated activity location x_i_, A_0_=<A_i_> is an average amplitude of the Ca^2+^ trace (<…> corresponds to averaging over all temporal samples), 𝜀=0.0001 * dt / 2πσ∼10^-7^ s/sm is a regularization parameter, and 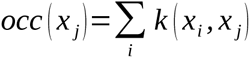 is the occupancy computed for each spatial bin x_j_ and summed over all time points i. The sampling rate was *fs=1/dt*=45 Hz.

To quantify the statistical significance of place fields, the spatial information (SI) (Skaggs et al., 1992) between location and Ca^2+^ activity has been determined according to the following formula:

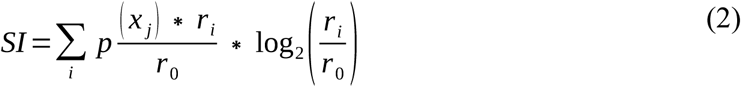

where j is the index of a spatial bin, p(x_j_) = 1 / (Σ_j_ occ(x_j_)) * occ(x_j_) is the normalized occupancy of spatial bin x_j_, r_i_ and r_0_ are the mean event rate in the bin and the overall mean event rate, respectively. The true value of SI has been compared to a shuffle distribution. To this end, in each of the 200 shuffles, the calcium trace was circularly shifted with respect to the corresponding trajectory by a random value ranging from 60 seconds to session duration minus 60 seconds. The cell was defined as a significant place cell (PC) if its original SI value exceeded the 95^th^ percentile of the shuffle SI distribution.

To compare the place field sizes and mean in-field activities, we extracted the contour line of the firing map at 0.3 of the maxima. The mean in-field activity was defined as the mean value inside this contour, while field size was defined as the within-field number of pixels multiplied by the pixel size (1.8-by-1.8 cm^2^). In cases with multiple detected contours, each was treated as an independent field. To obtain the shuffled distribution of PF sizes, place field detection was repeated on circularly shuffled rate maps, as described above.

For the split-half correlation (SHC), each calcium trace and the corresponding trajectory (at running speed larger than 5 cm/s) were split into two halves, and each half of the data yielded a corresponding rate map. Pearson correlation between place maps of the two halves was compared to a distribution obtained by circular shuffling as described above. A cell was determined as a significant SHC Place Cell (SHC PC) if the original SHC value was above the 95^th^ percentile of the shuffle SHC distribution.

### 2.4 Temporal analysis

Cell footprints were tracked according to an established routine (Sheintuch et al., 2017). In brief, for each session recorded from the same subject, the spatial footprints of all cells were projected onto a single image. Then, a field of view (FOV) from the session with the highest correlation to the other FOVs was chosen as the reference. Afterward, a series of translations and rotations was applied to the FOV of each session to maximize the cross-correlation with respect to the FOV of the reference session. That provided the locations of all spatial footprints within the reference coordinate system (Sheintuch et al., 2017). Next, the amount of spatial overlap between cell pairs and the centroid distance between them (similarity measures) were used to yield bimodal distributions of neighboring ROIs from the aligned projection. Each distribution was modeled as a weighted sum of unimodal distributions of the same cells (presumed to be the same cells active on different sessions) and different cells (presumed different cells potentially active in the same session) (Sheintuch et al., 2017). The fit errors from two similarity measures were compared to provide the optimal model. Corresponding optimal unimodal distributions were subjected to Bayesian probabilistic modeling to determine the probability of two neighbors being the same cell (identity probability). Lastly, those probabilities were used to initiate a clustering procedure. That procedure converged upon maximizing the within-cluster identity probabilities and minimizing the across-cluster identity probabilities (Sheintuch et al., 2017).

Before multi-session cell registration, the parameters of the multi-session alignment and probabilistic modeling algorithms were optimized in the following way. The radius of a circular area around each pixel in which optical flow was registered, and the threshold separating neighboring cells, were chosen by visual inspection of false-positive and false-negative error fractions. If a subset of sessions looked very different from another subset from the same animal (potentially reflecting optical lens reattachment or observing different outputs of the CNMF-E algorithm), the two subsets were treated as if coming from different subjects.

### 2.5 Place cell recurrence simulations

To simulate random place cell appearance (**Fig. 4K**), each tracked cell had a probability of being a place cell of 0.175, reflecting the total proportion of place cells detected in our study. For each day, each cell was treated independently, providing a simulated population of trackable simulated place cells with a uniform probability of being a place cell.

### 2.6 Correlation and population vector analysis

Four subsets of cells were used in this analysis, given the fixed time lag between the tracked sessions: all cells, non-place cells (NPCs, neurons that were determined as a place cell in neither of the tracked days), place cells on one of the tracked days (PC/NPC and NPC/PC), and place cells on both tracked days (PC/PC). For each pair of cells in each subpopulation, Pearson’s correlation coefficient between their corresponding place fields was determined.

To define stable cell pairs, we applied a threshold of 0.3 to Pearson’s correlation, since, by visual inspection, this value reflected the upper tail of the distribution of all cell correlations. Population vector correlation was calculated according to the following formula:

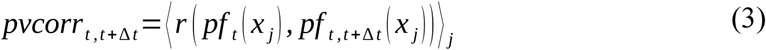

Where pf_τ_(x_j_) corresponds to a 1D vector being a stack of all place fields at position x_j_ in session t, r is Pearson’s correlation coefficient, and <…>_j_ indicates averaging across spatial pixels. Both ratemap and population correlation analysis were performed for the SI and the SHC PCs; cell pairs were split into ones tracked in consecutive and non-consecutive sessions.

### 2.7 Center-of-mass analysis

Fields were redetermined with a contour line at 0.6 of the maximum value of the rate map. The center of mass (CM) of a rate map pf(x_j_) within that contour was defined as

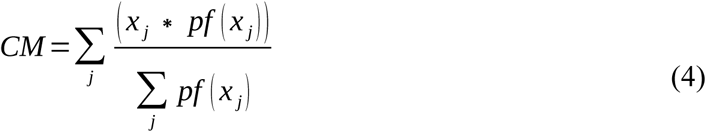

For each cell, the CM shift was defined as the distance between two CMs detected in each session. In the case of multiple detected CMs, CM shift was defined as the minimum of all values. For a pair of placefields, the change in PF size was determined as an absolute difference in the number of pixels within the PFs multiplied by bin size (1.8 cm)^2^. To fairly compare PC/NPC and PC/PC populations, which had very different magnitudes, ten random down-samples of the PC/NPC population were performed to match the size of the PC/PC populations, and the result was averaged across the subsamples. Changes of CM shifts and PF sizes over time were computed only on PC/PC pairs and compared against randomized control data. To this end, for each tracked PC on the first day, a random PC was picked from the pool of tracked PCs on the second day in order to compute random CM shifts and PF size changes. This procedure was repeated ten times and the results were averaged over the repetitions.

For rate map correlation- and population vector-analyses, field shifts were computed for consecutive and non-consecutive tracked cell pairs and SI and SHC PCs.

### 2.8 Positional decoding

In this analysis, we used deconvolved spike-like activity instead of calcium transients. For each session, the running trajectory was linearly interpolated to match the sampling frequency of the calcium activity. Both the cell activity and the trajectory were binned into 0.2 s intervals. All even-numbered bins were concatenated and used as training data, and all odd-numbered bins were concatenated and used as testing data. An epsilon-support vector regression (Smola and Schölkopf, 2004) with a Gaussian RBF kernel with 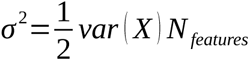 was applied to the training data to fit the neural activity 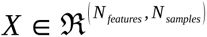 to the running trajectory. The testing data were then used to generate predictions, which were compared to the testing trajectory; the Euclidean distance between the true and predicted positions was taken as the decoding error.

The position decoding was performed using cell activity from three classes within a session: all cells, NPCs, and PCs. For the first two classes, cells were randomly subsampled five times to match the size of the corresponding PC population, and decoding errors were averaged across subsamples.

For each cell class, the analysis was repeated using shuffled data. To this end, the indices of the cells in the testing data were permuted, while the training data retained the original (non-permuted) cell indices.

This analysis was also performed for all subjects to compare two different PC detection methods with cells that were not identified by either of the methods (NPCs). Here, the cell populations were subsampled five times to match the size of a smaller PC population, and decoding errors were averaged across subsamples. In case the smallest population was less than 15 cells, the session was omitted from the analysis.

### 2.9 Computing correlations between cell loads and mean firing rates

Contribution of individual cell types to the population activity was measured as the correlation between principal component loads and firing rate. To fairly compare the PC and NPC populations, the NPC population was randomly downsampled ten times to match the size of the corresponding PC population, yielding the matrix *A^^^* (*t*) containing activities of equally sized populations of PCs and NPCs. Then, the time-dependent firing rate *FR(t)* was computed from *A^^^* (*t*) as follows:

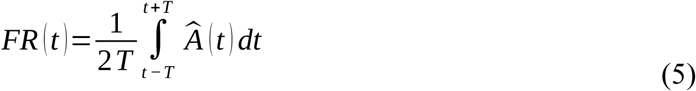

Where *T=12/fs*∼0.27s is half of the sliding window width and f*s*=45 Hz is the sampling rate of the transients.

The principal component loads were defined in the following way:

1. The cell activity covariance matrix *C(A^^^)* was computed.
2. *C(A^^^)* was subjected to eigenvalue decomposition, yielding the matrix: 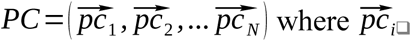 is a normalized eigenvector of *C(A^^^)* with an eigenvalue *λ_i_*. The eigenvectors in *PC* are sorted according to their eigenvalues.
3. The absolute loads of the first ten principal components (with the largest eigenvalues) were computed in the following way:

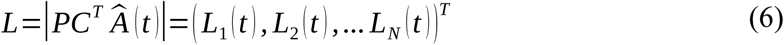

Each component load *L_i_(t)* was correlated with the mean firing rate *<FR* (*t*)*>,* where firing rate was averaged either over all PCs or subsampled NPCs and the results were averaged across ten subsamples. This analysis was done for both place cell detection methods. For pairs of sessions (cross-day analysis we used the matrix *Ã*(*t̃*) that contains transients of all tracked cells in a session pair with the populations of NPCs downsampled to match the size of the corresponding PC population. The data was pooled according to the temporal separation between the tracked sessions and the results were averaged across three principal components with the highest eigenvalues. Just like in the same-day analysis, the results were averaged across the subsamples.

## 3. Results

### 3.1 Two-dimensional place cells show higher activity levels and more defined place fields than non-place cells

We re-analyzed previously published recordings of dorsal CA1 principal neuron (PN) activity obtained by wide-field head-mounted optical imaging of Ca^2+^ transients, reported by GCaMP6f fluorescence, in freely moving mice (Chenani et al., 2022). In this dataset, animals explored a round arena in 15-minute daily sessions for several consecutive days. We restricted our analysis to sessions with sufficient exploratory behavior and Ca^2+^ recording quality (see **Section 2.2**), which included 10 mice and 10 consecutive daily recording sessions. We identified individual cells and extracted the corresponding Ca^2+^ fluorescence traces using standard preprocessing methods (CaImAn, **Fig. 1A-C** and **Methods**) (Giovannucci et al., 2019). We detected 324±203 cells per session/animal. The distribution of PC fields’ centroids over time varied across mice (**Supplementary Figure 1A**), and mice sampled the arena with a higher density towards the borders (**Supplementary Figure 1B**).

**Figure 1.**
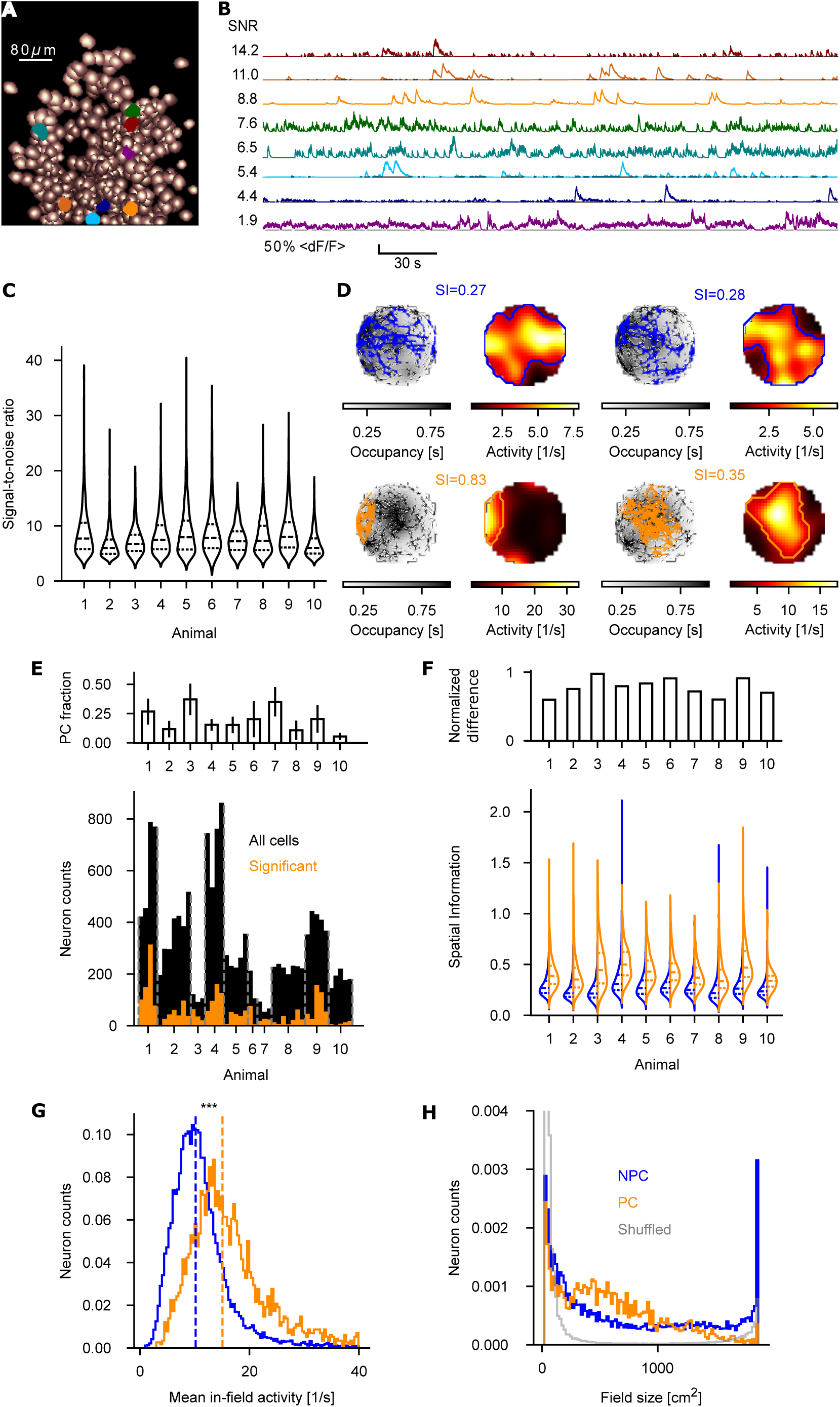
Place cells have a higher firing rate and a preferred place field size. **A**: Example field of view with footprints of cells detected by CaImAn in a 15-minute recording session. Pixel brightness indicates the correlation of each pixel with itself across all the other frames within the session. Colored footprints correspond to cells in **B**. **B**: Five-minute-long deconvolved calcium traces of 8 example cells, sorted by SNR values. **C**: Distributions of SNR values for 10 mice. **D**: Example PFs. Top: NPCs, blue; bottom: PCs, orange. Left: running trajectories (black) with calcium event coordinates (blue/orange). Right: normalized rate maps. Contour lines correspond to 30% maximum and indicate the place fields. **E**: Cell counts across mice and sessions (total 14590 cells). Each bar depicts a single session; individual animals are divided by vertical dotted lines. Black: total cells; orange: PCs. Inset: Average PC fraction in each animal. **F**: SI content of PCs and NPCs per mouse (0.39±0.08 mean for PCs, 0.25±0.03 mean for NPCs, 0.27±0.03 bits mean for all cells). Inset: difference between the mean SI values of PCs and NPCs normalized by the square root of the combined standard variance of both cell types for each mouse. **G**: Distributions of mean in-field activity pooled across all animals and sessions (15.1 1/s mean for PCs, 10.1 1/s mean for NPCs). Kruskal-Wallis test (***: p=0, H=2092, n_PC_=3237, n_NPC_=18357). **H**: Distribution of field sizes pooled across all mice and sessions (mean for PCs: 590±432 cm^2^; mean for NPCs: 753±631 cm^2^). Grey: control distribution obtained from randomization (see **Section 2.3**). **D, F, G, H**: Blue: NPCs; orange: PCs. **C, F**: Dotted lines indicate the median, upper, and lower quantiles.

All ensuing analyses, except the direct comparison in **Supplementary Figure 2** and decoding in **Supplementary Figure 5** was performed on Ca^2+^ transients. A subset of cells demonstrated high spatial tuning of their DF/F trace, defined by their SI exceeding the 95%-tile of a shuffle distribution (**Fig. 1D**) (Skaggs et al., 1992). We defined this neuronal subset as PCs. PCs constituted 17.5±13.1% of all recorded cells per session (**Fig. 1E**), showed SI values and mean in-field activity significantly higher than non-place cells (NPCs, **Fig. 1F** and **G**). Moreover, PCs exhibited a preferred place field (PF) size (590±432 cm^2^), as indicated by the mode of the distribution of PF sizes, while NPCs did not (**Fig. 1H**). The distribution of PF size, obtained by shuffling (see **Section 2.3**), resembled the distribution of NPC PF sizes.

To test for methodological biases, we repeated these analyses using the deconvolved “spike” activity traces - obtained by application of the Online Active Set method to Infer Spikes (OASIS) (Friedrich et al., 2017) in the CaImAn software - instead of the DF/F trace. As we found only small quantitative differences (**Supplementary Figure 2**), we decided to perform ensuing analyses based on the DF/F traces.

### 3.2 Place cell codes are more stable than non-place cell codes

We registered our data across time by using multisession Field of View alignment, probabilistic modeling, and cell clustering (Sheintuch et al., 2017) and tracked cells across sessions (**Fig. 2A**), with a high cell exclusivity score (86.8±9.1%), low false-positive (4.4±5.8%), and false-negative errors (19.7±9.9%). This enabled us to follow the same neurons and their activity patterns across time and thus investigate the stability of spatial codes at the single-cell and population levels at different time scales.

**Figure 2.**
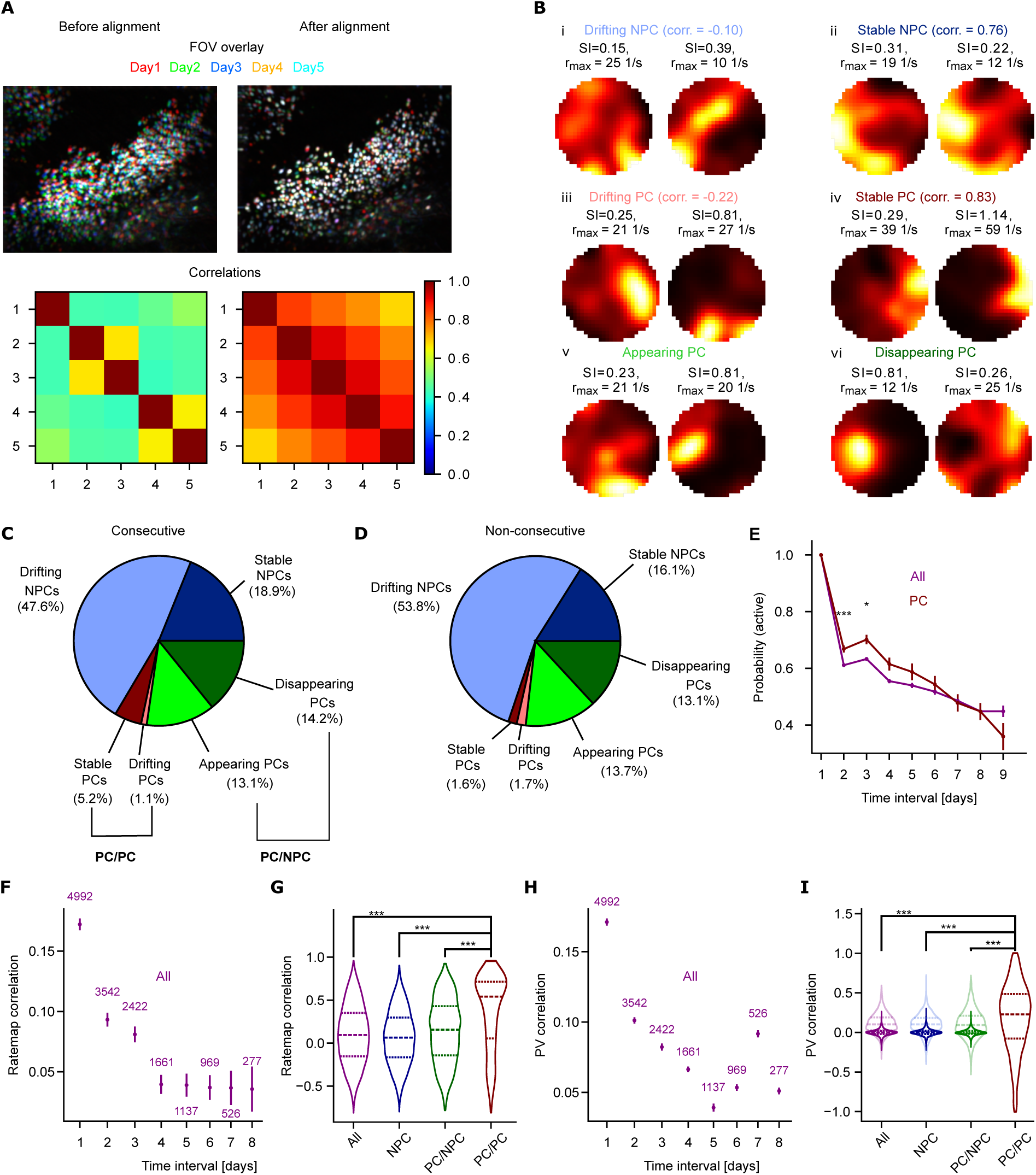
Place cells are more temporally stable at the single-cell and population level. **A**: Top: example overlay of cell footprints across five sessions, bottom: corresponding across-session FOV correlations of pixel intensity values. Right: before, left: after isometric alignment of the FOVs across sessions. **B:** Example cell pairs tracked across days. Drifting (i, light blue) and stable NPCs (ii, dark blue). Drifting (iii, light red) and stable PCs (iv, dark red), appearing (v, light green) and disappearing PCs (vi, dark green). **C:** Fractions of cell pairs tracked in consecutive sessions according to the classification in **B**. **D**: Same as C, but for non-consecutive session cell pairs. **E:** Recurrence probability of ROI for all cells and PCs. One-way ANOVA to compare all and PC recurrence probability on the same day basis (***: p=0.00020, statistic=13.8, n_all_=5597, n_pc_=767; *: p=0.013, statistic=6.2, n_all_=4366, n_pc_=470). **F**: Average rate map correlations decay with time. Numbers correspond to the number of cell pairs for each time point. **G**: Distributions of rate map correlations for cell pairs classified according to B. Kruskal-Wallis test (***: p∼10^-73^, H=327, n_all_=15526, n_pc/pc_=663; ***: p∼10^-88^, H=398, n_npc_=10676, n_pc/pc_=663; ***: p∼10^-50^, H=222, n_pc/npc_=4187, n_pc/pc_=663). Dotted lines are the median, upper, and lower quantiles, respectively. **H:** Population vector correlations decay with time. Numbers indicate the number of cell pairs for each time point. Note that the marked increase for the 7-day interval is putatively caused by a low number of cells tracked at this time interval. **I**: Distributions of population vector correlation for cell pairs classified according to **B**. Kruskal-Wallis test (***: p=0, H=2767, n_all_=12274, n_pc/pc_=11552; ***: p=0, H=2757, n_npc_=12274, n_pc/pc_=11552; ***: p=0, H=2667, n_pc/npc_=12274, n_pc/pc_=11552). Darker colors correspond to the values obtained by down-sampling each population size to the size of the PC/PC population, while lighter colors show the same results without the subsampling. **E, F, G, H, I**: Purple: all cells; blue: NPCs; green: PC/NPCs; red: PC/PCs. **E, F, H**: Error bars correspond to S.E.M.

We used the correlation of the firing map between consecutive days for PCs and NPCs as a metric of spatial coding stability. Hence, we categorized cell pairs with correlation values > 0.3 as stable and cell pairs with correlation values < 0.3 as drifting. We thus obtained the following six classes. Drifting and stable NPC pairs did not exhibit significant place tuning on either of the tracked days, and their corresponding rate map correlation value across days was < 0.3 and > 0.3, respectively (**Fig. 2B** i and ii). Drifting and stable PC pairs exhibited significant place tuning on both tracked days (PC/PC), and their corresponding rate map correlation value across days was < 0.3 and > 0.3, respectively (**Fig. 2B** iii and iv). Appearing and disappearing PC pairs exhibited significant place tuning either on the second but not on the first day, or on the first but not on the second day (NPC/PC and PC/NPC, respectively, **Fig. 2B** v and vi). When considering sessions on consecutive days, the majority of cell pairs (*c.a.* 50%) were drifting NPCs, the next more abundant class (*c.a.* 30%) were appearing and disappearing PCs (13% and 14% respectively). Stable NPCs comprised 19%, while stable PCs were only about 5%, and the remaining 1% were drifting PCs (**Fig. 2C**). Similar fractions were obtained for the cell pairs from non-consecutive sessions, with the only difference being that stable PCs were less prominent (less than 2%, **Fig. 2D**).

We then focused on the dynamics of spatial coding at longer time scales. For each cell, the probability of being active in any pair of two sessions decreased with elapsed time. This decay was very similar between all cells and PCs, with the notable exception of 2- and 3-day intervals, where PCs were significantly more likely to reappear than all cells (**Fig. 2E**). In line with these results, also the rate map correlations of all cells decreased as a function of time (**Fig. 2F, Supplementary Figure 3**). Rate map correlations in the PC/PC cells, however, were significantly higher than in the other neuronal subpopulations (**Fig. 2G**). We then investigated the spatial information contained in the covariance structure of the population activity by using population vector (PV) correlations. Also, in this case, the PV correlations of PC/PC pairs were significantly higher than those of other neuronal subpopulations (**Fig. 2H, I**, light colors). To account for different numbers of neurons in the individual subpopulations, we repeated the PV correlation analysis while down-sampling the size of NPCs and all cell populations, again yielding highly consistent outcomes (**Fig. 2I**).

Overall, even though PCs are a small fraction of PNs in CA1, the neural codes of PCs are less susceptible to drift as compared to the codes of NPCs.

### 3.3 Place fields of place cells shift less than those of non-place cells

We then investigated the temporal stability of PFs using single-cell level measurements. To this end, we estimated the center of mass (CM) of all PFs (PC/NPCs and PC/PCs) and calculated their shift in time, i.e. the distance between the CMs of PFs of the same cell at two different time points (**Fig. 3A**). To account for different numbers of neurons in the individual subpopulation, we also performed ten random down-samples of the PC/NPC population to match the size of the PC/PC population.

**Figure 3.**
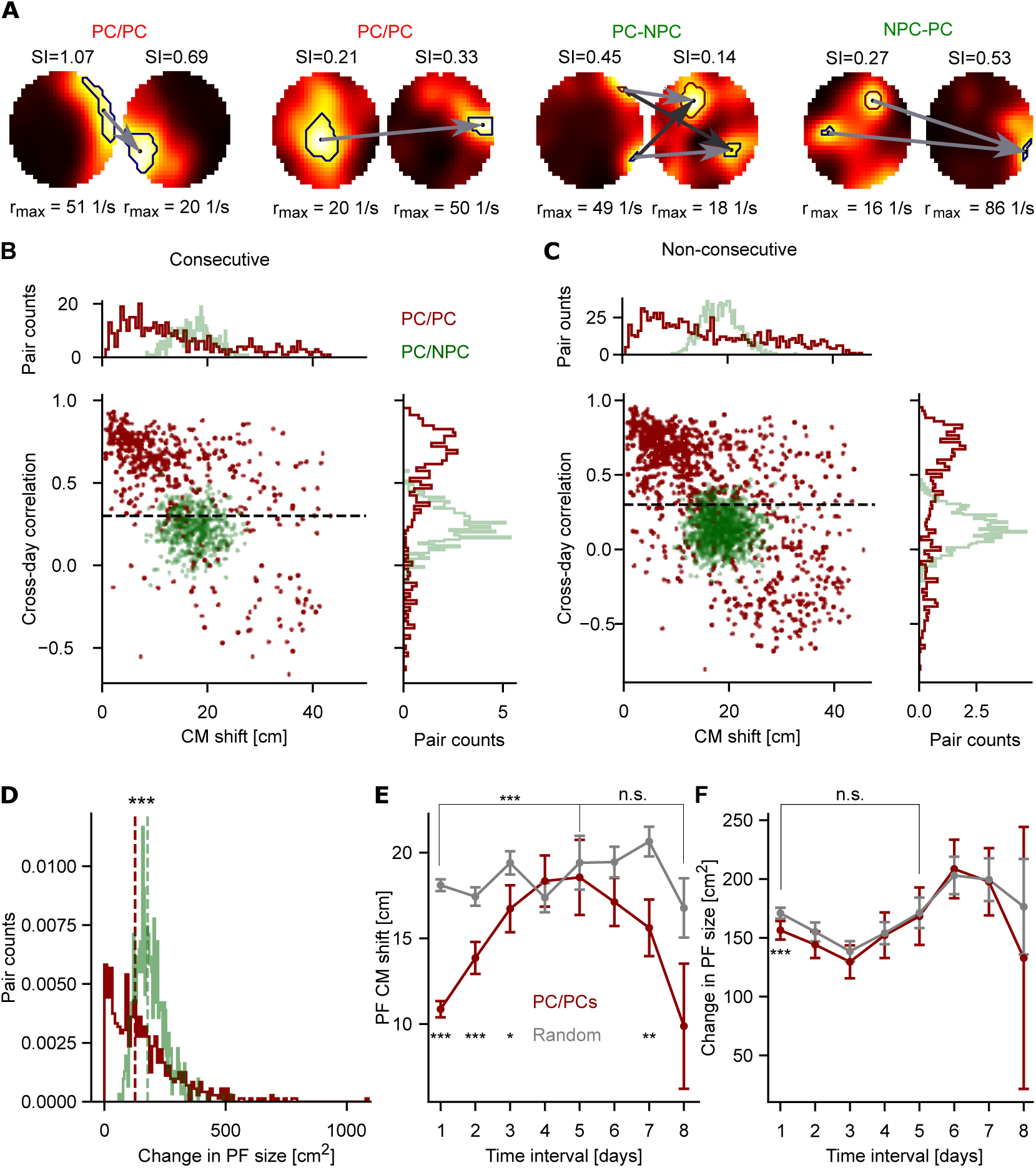
Reduced drift and field size change of place cells. **A:** Example place field shifts of two PC/PC pairs (far left, left), a disappearing PC pair (PC-NPC, right), and an appearing PC pair (NPC-PC, far right). In each ratemap, contours indicate 0.6 of the map’s maximum value (see **Section 2.7**). Arrows (grey for a single drifting field, additional dark grey for pairs of shifting fields) connect the center of mass within each contour and indicate all possible pairs of field shifts. In case of multiple contours detected, only the smallest shift value is considered for each cell pair. **B:** Joint distribution of cross-day rate map correlation vs. CM shift for down-sampled PC/NPC and PC/PC populations, tracked for consecutive sessions (ten random down-samples of the PC/NPC population were performed to match the size of the PC/PC population). Dots are cell pairs; the dashed line depicts the stability threshold of 0.3 introduced in Fig. 2. Top: marginal distribution of CM shifts; right: marginal distribution of cross-day correlations. **C**: Same as in **B**, but for the non-consecutive sessions. **D**: Distribution of changes in PF size for the PC/PC and down-sampled PC/NPC populations. Kruskal-Wallis test (***: p∼10^-23^, H=99, n_pc/npc_=663, n_pc/pc_=663). **E**: Average CM shift as a function of temporal separation for all trackable PC/PC pairs (red), compared to a random control (grey, see **Section 2.7**). Linear regression applied to the PC/PC data either during the first five days interval or during the last three days interval (***: p_1-5_∼10^-10^, F=44, df=1, n.s.: p_6-8_=0.09, F=2.8, df=1), Kruskal-Wallis test comparing the PC/PC data to a random control on a daily basis (***: p_1_∼10^-38^, H=167, n_pc/pc_=311, n_rand_=311, ***: p_2_∼10^-5^, H=18, n_pc/pc_=117, n_rand_=117, *: p_3_=0.023, H=5, n_pc/pc_=76, n_rand_=76, **: p_7_=0.005, H=7.9, n_pc/pc_=34, n_rand_=34). **F**: Same as in **E**, but for the change in the place field size. Linear regression applied to the PC/PC data either during the first five days interval or during the last three days interval (n.s.: p=0.29, F=1.1, df=1, n.s.: p=0.097, F=2.83.3, df=1),Kruskal-Wallis test comparing the PC/PC data to a random control on a daily basis (***: p _1_∼10^-5^, H=20, n_pc/pc_=311, n_rand_=311) **A, B, C, D**: Green: PC/NPCs; red: PC/PCs. **E, F**: Error bars correspond to S.E.M.

As expected, PC/PCs neuronal pairs with higher rate map correlations also displayed lower CM shifts, whereas neurons with lower rate map correlations demonstrated higher CM shifts. This resulted in an overall inverse dependency between rate map correlations and CM shifts for PC/PCs. For PC/NPCs, the CM shifts and correlations were unimodally distributed around chance levels of Pearson’s r and a CM shift of about 10-25 cm. Those results were consistent for PC/PC pairs detected on consecutive (**Fig. 3B**) and non-consecutive (**Fig. 3C**) sessions. In addition, PC/PC neuronal pairs showed size changes significantly smaller than PC/NPCs (**Fig. 3D**), thus demonstrating consistent PF structure in CA1 for stable PCs. For up to a five-day interval between the sessions, the PF CM shifts demonstrated a significant increase, whereas shifts after day 5 no longer showed a significant trend (**Fig. 3E**). Up to a three-day interval, the shifts were significantly lower than those of randomized control data. Moreover, PFs did not demonstrate any significant difference in the change of PF size as compared to randomized data except for one-day intervals (**Fig. 3F**), where PF sizes change slightly less than in random control data. These results confirm that active PCs with consistent spatial tuning experience a lower degree of representational drift, mostly correlating to CM shift but not PF size changes.

### 3.4 An alternative method for determining place cells provides a similar number of place cells with different spatial tuning

An alternative method to identify place cells does not rely on SI, but rather relies on the correlation between the two place maps derived from the first and the second halves of a recording session to exceed the 95^th^ percentile of a distribution of such correlations obtained from calcium trace randomization (Keinath et al., 2022) (see **Section 2.3**). This method is thus referred to as Split-Half Correlation (SHC). Importantly, it was reported to reveal an alternative and only partially overlapping population of PCs (Grijseels et al., 2021). We thus wondered whether using the SHC definition for PCs would affect the basic properties of PCs and NPCs as compared to SI-defined place cells. To test the applicability of the SHC approach for our data, we quantified the effect of transient bleaching within the sessions by computing the mean fluorescence in the first and the second halves of each recording. Although fluorescence values were significantly lower in the 2nd halves of the sessions (**Supplementary Figure 2E,** Wilcoxon rank-sum test), the observed difference was only 4.4% and should only have a minor effect on our results.

For our data set, both detection methods yielded similar PC fractions in each session and subject (SI-PCs: 17.5±13.1% and SHC-PCs: 17.0±11.5% of all recorded cells, **Fig. 4A**, **B**), but the populations of the two different kinds of PCs had only a limited overlap (**Fig. 4C)**. The amounts of SI in SI-PCs and SHC-PCs were similar (**Fig. 4D**) and the normalized SHC difference between SHC-PCs and SHC-NPCs (1.98±0.09, **Fig. 4E**) was more than twice as high as the normalized difference in SI between SI-PCs and SI-NPCs (**Fig. 1F**, 0.78±0.12). This suggests that the use of SHC could provide a clearer distinction between the PCs and NPCs, yet at the cost of inconclusive SI.

As expected, PCs detected only by the SI method (unique SI-PCs) showed higher SI content than PCs detected only by the SHC method (unique SHC-PCs), and unique SHC-PCs showed higher SHC value than unique SI-PCs. However, PCs detected by both methods (overlapping PCs) had higher SI content and a higher SHC value than the corresponding unique PCs (**Fig. 4F**, **G, H**). In addition, unique SHC-PCs exhibited a broader PF size than unique SI-PCs, while overlapping PCs had PF widths similar to SI-PCs (**Fig. 4I**).

Despite these differences, both detection methods yielded a similar decrease in cell recurrence probability (**Fig. 4J**). For both methods, the number of PCs tracked over time decreased as a function of time (**Fig. 4K**). Interestingly, SI-PCs were significantly more likely to recur as PCs compared to both SHC-PCs and PCs from a simulation of surrogate PC counts in which cells are randomly assigned to be a PC with a uniform probability (**Fig. 4L**, see **Section 2.5**). We therefore decided to compare the temporal stability of PCs and NPCs detected using both detection methods.

**Figure 4.**
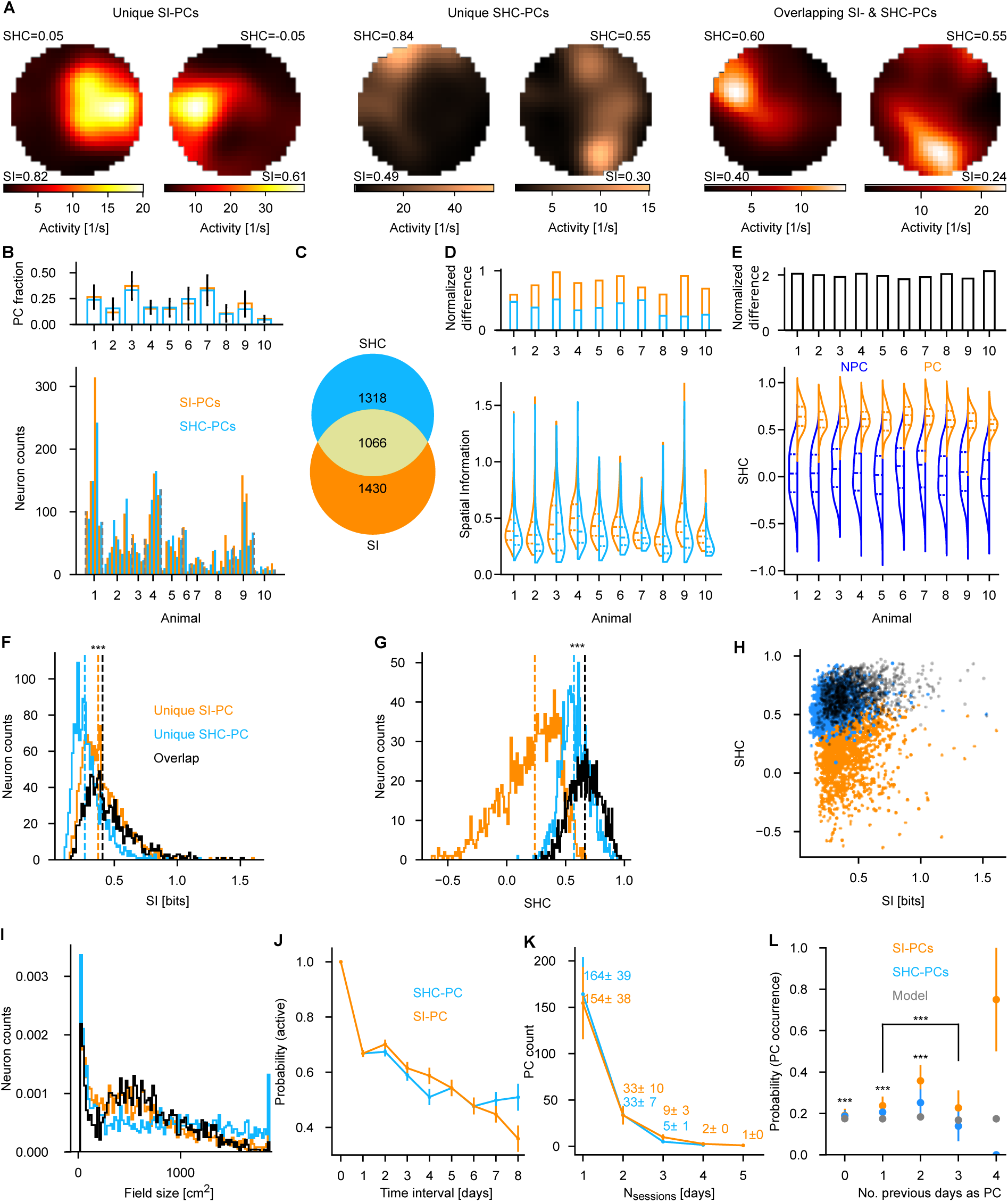
Overlap between the place cell detection methods. **A**: Example place fields of unique SI-PCs (left), unique SHC-PCs (center), and overlapping SI- and SHC-PCs (right). **B:** Comparison between the within-session counts of SI-PCs (orange) and SHC-PCs (light blue). Inset: mean count of SHC PCs per subject. **C:** Relative sizes as well as the overlap of SI-PC and SHC-PC populations. **D**: SI content of SI-PCs and SHC-PCs. Inset: normalized difference between each distribution for each animal. **E:** Distribution of the split-half correlations of SHC-NPCs (blue) and SHC-PCs (orange) per animal. Inset: difference between the split-half correlation values of SHC-PCs and SHC-NPCs normalized by the combined standard deviation of both cell types within each mouse (see Fig. 1F for comparison). **F**: SI content of unique SI-PCs, unique SHC-PCs, and overlapping PCs. Kruskal-Wallis test (***: p∼10^-10^, H=40, n_si-unique_=1430, n_overlap_=1066) **G**: SHC value of unique SI-PCs, unique SHC-PCs, and overlapping PCs. Kruskal-Wallis test (***: p∼10^-58^, H=257, n_shc-unique_=1316, n_overlap_=1066). **H**: Scatter plot of SI content vs. SHC value for unique SI-PCs (orange), unique SHC-PCs (blue), and overlapping PCs (black). I: Distribution of field sizes of unique SI-PCs (595±453 cm^2^, **see** Fig. 1H), unique SHC-PCs (786±575 cm^2^), and overlapping PCs (630±408 cm^2^). **J**: Recurrence probability of ROI for SI- and SHC-PCs. **K**: Histograms of days for which a tracked ROI was an SI- or an SHC-PC. Error bars indicate the count per animal. **L**: Probability of PC formation as a function of the number of days in which the PC was significant. Two-way ANOVA to compare the recurrence probability of SI-PCs to the one of SHC-PCs (***: p_meth1-3_∼10^-8^, F_meth1-3=_28.7, df_meth1-3_=1, p_time1-3_=0.29, F_time1-3=_1.24, df_time1-3_=3, p_intersect1-3_=0.11, F_intersect1-3=_2.03, df_intersect1-3_=3), binomial test to compare the recurrence probability of SI-PCs to the one of modelled cells as well as the recurrence probability of SHC-PCs to the one of modelled cells on the daily basis (***: p_si_∼10^-8^, statistic_si0_=0.16, p_si1_∼10^-14^, statistic_si1_=0.25, p_si2_∼10^-8^, statistic_si2_=0.31, p_shc0_∼10^-6^, statistic_shc0_=0.16). **B, C, D, F, G, H, I, J, K, L**: Orange: SI-PCs; light blue: SHC-PCs. **J, K, L**: Error bars correspond to S.E.M.

### 3.5 Place cells determined by the SHC significance are less stable than place cells determined by the SI significance

We again used the correlation of the firing map between consecutive days for SHC-PCs and SHC-NPCs as a metric of spatial coding stability and categorized cell pairs with correlation values > 0.3 as stable and cell pairs with correlation values < 0.3 as drifting. While the fraction of consecutive stable NPCs was significantly higher for the SHC method (SI-PCs: approx. 30%, SHC-PCs: approx. 35%, **Fig. 5A**), the stable fractions increased for consecutive PC/NPCs (approx. 50%, **Fig. 5B**). The gap between the proportions of cells identified by the two methods became more vivid for PC/PC pairs (approx. 80% SI cell pairs vs. approx. 60% SHC pairs for consecutive cell pairs, **Fig. 5C**). Similar trends were observed in non-consecutive cell pairs, but the corresponding proportions were lower (NPCs: approx. 20%, **Fig. 5D**, PC/NPCs: approx. 30%, **Fig. 5E**, approx. 50% SI-PC/PCs vs. approx. 40% SHC-PC/PCs, **Fig. 5F**). We next compared the dynamics of spatial coding at longer time scales for SI- and SHC-detection methods, and found that the stability of fields - in terms of rate map correlation - decreased as a function of time, and PCs were more stable than NPCs (**Supplementary Figure 4A**, see **Fig. 2F** and **Supplementary Figure 3A**) for both methods. However, SI-PCs were significantly more stable than SHC-PCs (**Supplementary Figure 4A**). On the contrary, when we analyzed PV correlations, SI-PCs were significantly more stable than SHC-PCs, depending on the time difference between the sessions (**Supplementary Figure 4B**). Moreover, the SI method provided significantly higher ratemap and PV correlations for pooled data (**Fig. 5G, H**).

**Figure 5.**
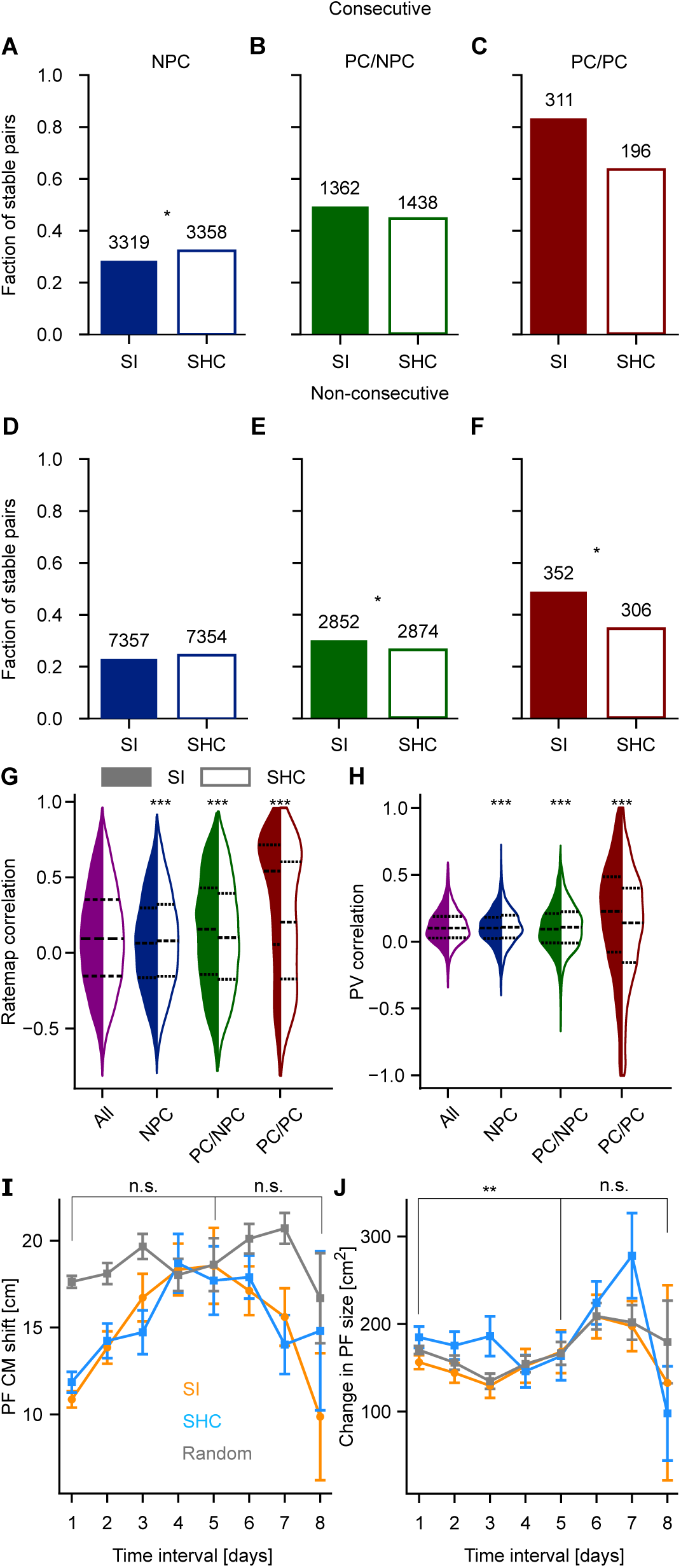
Place cells detected by using the split-half correlation method appear less stable than place cells detected by using the spatial information method. **A, B, C**: Fractions of stable (correlation > 0.3) cell pairs (NPC, blue, **A**; PC / NPC, green, **B**, and PC / PC, red, **C**) tracked in consecutive sessions for both detection methods (solid bars = SI, empty bars = SHC). Chi-squared test (p_npc_=0.013, statistic=6.2, df=1; p_pc/npc_=0.12, statistic=2.4, df=1; p_pc/pc_=0.07, statistic=3.4, df=1). Numbers above each bar are the total number of cell pairs. **D, E, F**: Fractions of stable (correlation > 0.3) cell pairs (NPC, blue, **D**; PC / NPC, green, **E**, and PC / PC, red, **F**) tracked in non-consecutive sessions for both detection methods (solid bars = SI, empty bars = SHC). Chi-squared test (p_npc_=0.12, statistic=2.5, df=1; p_pc/npc_=0.022, statistic=5.2, df=1; p_pc/pc_=0.021, statistic=5.3, df=1). Numbers above each bar are the total number of cell pairs. **G**: Ratemap correlation pooled across all tracked sessions for both PC detection methods (solid-SI, empty-SHC methods, respectively). Kruskal-Wallis test (***: p_npc_=0.00056, H_npc_=12, n_si-npc_=10676, n_shc-npc_=10712; ***: p_pc/npc_∼10^-5^, H_pc/npc_=17, n_si-pc/npc_=4187, n_shc-pc/npc_=4312; ***: p_pc/pc_∼10^-11^, H_pc/pc_=44, n_si-pc/pc_=663, n_shc-pc/pc_=502). **H**: Same as in **G**, but for the population vector correlation. Kruskal-Wallis test (***: p_npc_∼10^-7^, H_npc_=26, n_si-npc-pix_=13357, n_shc-npc-pix_=13357; ***: p_pc/npc_=0.00018, H_pc/npc_=14, n_si-pc/npc-pix_=13357, n_shc-pc/npc-pix_=13357; ***: p_pc/pc_∼10^-52^, H_pc/pc_=232, n_si-pc/pc-pix_=11552, n_shc-pc/pc_=11191). **I**: Average CM shift as a function of temporal separation for all trackable SI- and SHC-PC/PC pairs compared to a random control (gray, see **Section 2.7**). Two-way ANOVA to compare the SI- and the SHC-PC/PC data (n.s.: p_method_=0.57, F_method_=0.3, df_method_=1; p_time_∼10^-13^, F_time_=21, df_time_=3; p_intersect_=0.43, F_intersect_=0.9, df_intersect_=3, n.s.: p_method_=0.0.89, F_method_=0.018, df_method_=1; p_time_=0.12, F_time_=2, df_time_=3; p_intersect_=0.0.74, F_intersect_=0.4, df_intersect_=3). **J**: Same as in **I**, but for the change in PF size. Two-way ANOVA to compare the SI- and the SHC-PC/PC data (**: p_method_=0.0022, F_method_=9.4, df_method_=1; p_time_=0.49, F_time_=0.8, df_time_=3; p_intersect_=0.46, F_intersect_=0.85, df_intersect_=3, n.s.: p_method_=0.25, F_method_=1.3, df_method_=1; p_time_=0.07, F_time_=2.3, df_time_=3; p_intersect_=0.53, F_intersect_=0.7, df_intersect_=3). **A, B, C, D, E, F, G, H**: Purple: all cells; blue: NPCs; green: PC/NPCs; red: PC/PCs. **I, J**: Orange: SI-PCs; light blue: SHC-PCs. Error bars correspond to S.E.M.

Overall, the SI method identifies neurons with higher session-to-session stability than the SHC method. Hence, by comparison, the SHC method leads to faster drift than the SI method.

We then investigated whether the choice of SI or SHC as a PC detection method would affect the properties of PCs over time. The general trends observed for the cross-day ratemap correlations, CM shifts, and changes in the PF size (**Fig. 3**) were also prominent for SHC-PC/PCs (**Supplementary Figure 4C-E**). Note that the data for 7 and 8-day intervals between tracked sessions were very sparse in comparison to session pairs with a smaller separation (**Supplementary Figure 4F, G**, see **Fig. 2H, 3E-F, 5I-J**), limiting the interpretability of the analysis outcomes for these intervals. Comparison of CM shift over sessions did not yield any significant difference between the SI-PC/PCs and SHC-PC/PCs (**Fig. 5I**). However, comparison of PF sizes over sessions, showed a lower change in PF size up to a 5-day interval when using the SI detection method (**Fig. 5J**). Altogether, this indicates that the SI method leads to the identification of neurons with sharper PFs that are more stable through time when compared to the SHC method.

Thus, while SI- and SHC-based methods yielded a similar fraction of PCs, SHC-PCs tend to be less stable than SI-PCs and therefore show a higher rate of representational drift. Hence, the method used to identify PCs crucially influences the quantification of representational drift if only the subset of “place-selective” cells is considered.

While representational drift is generally considered a population phenomenon, we also sought to address the contribution of individual cell classes such as SI/SHC-PC/NPC. We thus first trained kernel support vector classifiers (see **Section 2.8**) to predict the 2D location of the mouse from spike data at each single time point. Training and test data were taken from the same session (see **Section 2.8**). Owing to the limited spatial coverage of the arena test decoding errors were generally above 10 cm but still significantly lower than shuffle controls (**Fig. 6A**). Decoding errors obtained by using only the individual subclasses of cells were almost identical. While decoding error was slightly higher for NPCs (**Fig. 6B**), it was basically identical for both PC detection methods, showing that PCs and NPCs show similar contributions to the population code within a session, irrespective of the detection method. However, the low number of stable place cells over days prevented evaluating the population-level drift using the decoder. We therefore quantified the correlations of mean firing rates of SI/SHC-PCs and respective NPCs with population patterns of largest variance (Principal Components, see **Section 2.9**; **Fig 6C, D**). If computing the principal components from the activity of a single session, the most prominent population patterns, *i.e.*, the absolute loads of the first three principal components, correlated more strongly with the PC firing rates than the NPC firing rates. This difference was significant for SI-PCs but not for the SHC-PCs and SI-PCs were significantly more correlated than SHC-PCs to the first principal component (**Fig. 6E**) indicating that SI-PCs contribute most to the prominent population patterns. To assess the stability of patterns over days, we computed the principal components from paired sessions with a certain time separation and pooled correlation coefficients between mean firing rates and loads of the first three principal components. Again correlations of PCs were higher than those for NPCs with the largest difference for SI-PCs (**Fig. 6F**). Correlations between PCs and cell type-specific firing rates were almost identical for intervals of up to four days, indicating stability of the contribution of PCs to the population pattern. For longer time differences (7 & 8 day intervals), PCs no longer made up for the largest variance, indicating inconsistent rate maps of PCs over these intervals owing to drift (see **Supplementary Figure 4F, G**). Thus, place field detection has an influence on the stability and drift of PCs, both on the single cell and the population code of space.

**Figure 6.**
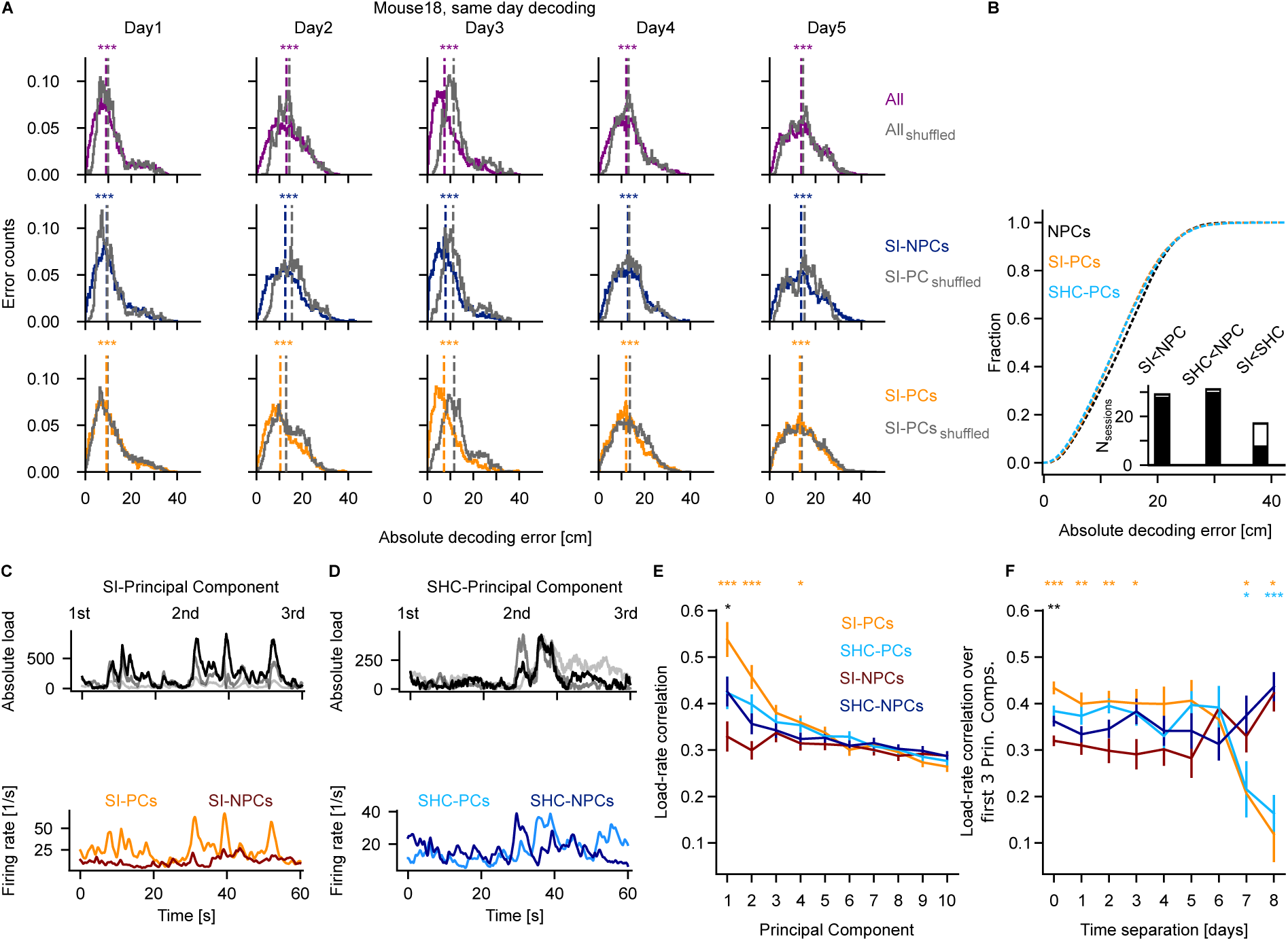
Comparison of population measures of SI and SHC PCs. **A**: Distributions of mean decoding errors for five sessions in one example animal (purple: all cells, dark blue: SI-NPCs, orange: SI-PCs against shuffled data (grey), see **Section 2.8**). Dotted lines highlight the medians of the distributions. Kruskal-Wallis test (Day1: p_all_∼10^-84^, H=380, n_all_=56, p_si-npc_∼10^-30^, H=134, n_si-npc_=56, p_si-pc_∼10^-15^, H=64, n_si-pc_=56; Day2: p_all_∼10^-147^, H=669, n_all_=22, p_si-npc_=0, H=1873, n_si-npc_=22, p_si-pc_∼10^-227^, H=1035, n_si-pc_=22; Day3: p_all_=0, H=4962, n_all_=169, p_si-npc_=0, H=4220, n_si-pc_=169, p_si-pc_=0, H=4989, n_si-pc_=169; Day4: p_all_∼10^-65^, H=293, n_all_=145, p_si-npc_∼10^-18^, H=78, n_si-pc_=145, p_si-pc_∼10^-78^, H=353, n_si-pc_=145; Day5: p_all_∼10^-38^, H=167, n_all_=55, p_si-npc_∼10^-34^, H=149 n_si-pc_=55, p_si-pc_∼10^-27^, H=119, n_si-npc_=55). **B**: CDFs of same-day decoding errors of SI-PCs (orange), SHC-PCs (light blue), NPCs (black, see **Section 2.8**), pooled over all sessions and animals. Inset: Number of sessions in which either SI-PCs or SHC-PCs yielded significantly lower decoding errors than NPCs or in direct comparison (black). The white bar indicates the total number of sessions with more than 15 cells per category. **C**: Top: the absolute loads of three principal components, derived from the activity of combined populations of SI-PCs and downsampled SI-NPCs in an example session, plotted over 60 seconds. Bottom: corresponding firing rates of SI-PCs and SHC-PCs. **D**: Same as in **C**, but for SHC-PCs/NPCs for the same time interval as in **C**. **E**: Correlation between the mean firing rate of SI-PCs (orange), SI-NPCs (dark red), SHC-PCs (light blue), and SHC-NPCs (dark blue) within sessions and loads of ten principal components sorted by their corresponding eigenvalues (see **Section 2.9**). One-way ANOVA to compare the correlations between three cell groups: SI-PCs vs. SI-NPCs (orange), SHC-PCs vs. SHC-NPCs (light blue), and SI-PCs vs. SHC-PCs (black) per principal component (p_si-pc/si-npc_∼10^-4^, statistic=17.8, n_si-pc_=37, n_si-npc_=37, p_si-pc/si-npc_∼10^-5^, statistic=24, n_si-pc_=37, n_si-npc_=37, p_si-pc/shc-pc_=0.042, statistic=4.3, n_si-pc_=37, n_si-npc_=37, p_si-pc/shc-pc_=0.0028, statistic=5, n_si-pc_=37, n_shc-pc_=41). **F**: Similar to **D**, but using cells tracked in a pair of sessions; the data is pooled across the first three principal components with the highest eigenvalues (see **Section 2.9**). The data for the time separation of zero corresponds to the data from **D** for the first three principal components. One-way ANOVA to compare the correlations between three cell groups: SI-PCs vs. SI-NPCs (orange), SHC-PCs vs. SHC-NPCs (light blue), and SI-PCs vs. SHC-PCs (black) per time separation between the sessions (p_si-pc/si-npc_∼10^-9^, statistic=40.5, n_si-pc_=148, n_si-npc_=148, p_si-pc/si-npc_=0.005, statistic=8.2, n_si-pc_=54, n_si-npc_=54, p_si-pc/si-npc_=0.0028, statistic=9.5, n_si-pc_=39, n_si-npc_=39, p_si-pc/si-npc_=0.018, statistic=6, n_si-pc_=27, n_si-npc_=27, p_si-pc/si-npc_=0.04, statistic=5, n_si-pc_=9, n_si-npc_=9, p_si-pc/si-npc_=0.013, statistic=17.9, n_si-pc_=3, n_si-npc_=3, p_shc-pc/hc-npc_=0.048, statistic=4.6, n_shc-pc_=9, n_shc-npc_=9, p_shc-pc/hc-npc_=0.0003, statistic=29, n_shc-pc_=6, n_shc-npc_=6, p_si-pc/hc-npc_=0.006, statistic=7.7, n_si-pc_=148, n_shc-pc_=164).

## 4. Discussion

We investigated the dataset generated by Chenani and colleagues by imaging dCA1 PN in mice freely foraging in an open arena (Chenani et al., 2022). Neural activity was recorded using wide-field head-mounted optical imaging of Ca^2+^ transients. We first defined PCs by using the significance of SI content and found higher temporal stability of PFs in PCs compared to NPCs, thus resulting in slower representational drift in PCs versus NPCs. We then detected PCs using the significance of the SHC value and found that while both the SI and SHC methods yield comparable numbers of PCs, the overlap between SI-PC and SHC-PC populations was only partial. Moreover, the firing rate maps of the independent PC subpopulations exhibited drift at different rates, with SI-PCs being more stable than SHC-PCs.

### 4.1 Place cells exhibit temporal stability higher than non-place cells

Optical imaging studies have consistently reported that PCs maintain spatial firing fields more stably over periods from days to weeks, whereas NPCs exhibit more labile and context-dependent activity (Dombeck et al., 2010; Ziv et al., 2013). Moreover, the probability of a PC re-occurring at a future time point has been shown to increase with the number of times that PC had already occurred in the past (Vaidya et al., 2025). Our results are consistent with these studies and expand the findings to mice moving freely in an open field. The higher temporal stability of the spatial codes of cells with higher spatial information (PCs) also corroborates a recent study reporting a positive relationship between contextual coding and long-term stability of coding properties (Keinath et al., 2022).

### 4.2 The choice of method to identify place cells affects the number and the characteristics of the cells detected

SI- and SHC-based PC detection methods have been applied to Ca^2+^ imaging data in several previous studies (Geva et al., 2023; Dombeck et al., 2010; Geiller et al., 2017; Grijseels et al., 2021). Keinath *et al*. defined PCs based on either SI or SHC in CA1 Ca^2+^ wide field imaging data obtained from mice freely navigating a morphing arena, and found a higher proportion of SHC-PCs than of SI-PCs (Keinath et al., 2022). A further study (Stefanini et al., 2020) - also focusing on CA1 Ca^2+^ wide field imaging data obtained from mice navigating an arena - demonstrated an unexpectedly low number of PCs. Moreover, a recent overview (Talpir et al., 2025) across studies suggests that Ca^2+^ imaging produces fewer PCs than electrophysiological recordings due to the lower sensitivity of Ca^2+^ indicators. Finally, significant differences between SI, SHC, and other methods for PC detection were also observed in synthetic datasets (Grijseels et al., 2021). The discrepancies between these findings challenge the robustness of classifying hippocampal neurons into PC and NPC based on Ca^2+^ imaging. By analyzing the same dataset obtained from mice freely navigating an arena with SI- and SHC-based PC detection methods, our study not only shows the stark differences between the two methods, but also that different datasets analyzed with the two methods can yield significantly different results (similar numbers of SI and SHC-PCs in our case versus different numbers in Keinath et al., 2022), thus adding to the above-mentioned variability.

### 4.3 Applying multiple place cell detection methods might lead to more accurate place cell estimation

Applying multiple independent significance criteria and designating only cells that meet all such criteria as PCs could decrease the variability in PC detection. As different detection methods lead to the identification of cell populations with a variable amount of overlap, the number of PCs identified by multiple methods is poised to be smaller than the number of PCs identified by a single method. Consistently, in our work, the number of overlapping SI- and SHC- PCs amounted to about 40% of the PCs identified either with SI or with SHC. This, however, greatly limits the statistical power of the analyses that can be carried out on the dataset (Grijseels et al., 2021) and, therefore, might be relevant only to study neuronal properties that highly correlate with a specific spatial location. Notably, this effect is strongly dependent on the threshold of detection for Ca^2+^ signals. In fact, using the SI and SHC significances simultaneously (Grijseels et al., 2021; Zong et al., 2022), together with setting thresholds for the PF size and the preferred mean within-field activity in two-photon optical recordings of Ca^2+^ transients obtained in an open arena, can lead to 48% PCs (Zong et al., 2022). This demonstrates that multi-criteria detection may be a valuable option when using two-photon imaging due to its higher sensitivity to neuronal activity.

### 4.4 The choice of method to identify place cells affects the quantification of representational drift

Representational drift in the mammalian brain refers to the gradual and ongoing changes in neuronal activity patterns that encode specific information - such as sensory inputs, learned tasks, or spatial location - even when behavior remains stable and consistent (Ziv et al., 2013; Mankin et al., 2012; Driscoll et al., 2022; Hainmueller and Bartos, 2018). Previous research examined representational drift using Ca^2+^ imaging in the dorsal hippocampus of mice navigating linear tracks or open fields (Ziv et al., 2013; Khatib et al., 2023; Dong et al., 2021; Rubin et al., 2015; Kinsky et al., 2018; Hayashi et al., 2023). However, these works did not explore how the PC detection method - often SI - affects drift quantification. Our findings that different detection methods identify distinct PC subpopulations - each exhibiting similar but not identical drift behaviors - underscore the importance of PC detection. This is particularly relevant when combining wide-field Ca^2+^ imaging - whose sensitivity is lower in comparison to two-photon imaging - and open-field (2D) foraging tasks - which produce much more non-uniform spatial sampling when compared to linear track (1D) spatial navigation.

### 4.5 Population coding approaches as an alternative to study representational drift

Early defining studies of PCs in the hippocampus (O’Keefe et al., 1978; O’Keefe and Dostrovsky, 1971) were limited by the small number of cells that could be recorded with available methods in each animal. This biased the field towards the analysis of single cells when investigating the properties of the hippocampal neural code. That, however, disregards the idea that non-place cells could also contain information about the animal’s position (Stefanini et al., 2020; Talpir et al., 2025) or that a specific location could be encoded by correlated activity between different neurons (Hazon et al., 2022; Nardin et al., 2023) rather than the activity of single neurons. In addition, several works have demonstrated a positive relationship between CA1 spatial coding and theta phase precession (McClain et al., 2019; Sloin et al., 2022). Conversely, hippocampal cells also encode other variables besides position, such as velocity (Stefanini et al., 2020; Góis and Tort, 2018; Iwase et al., 2020) or head direction (Acharya et al., 2016; Mankin et al., 2019; Yiu et al., 2022), which is particularly important in open arenas. These complex features of spatial codes are not easily captured by single-cell level analysis focused on position, but they can be investigated at the scale of populations by means of neural manifolds (Levy et al., 2023; Sun et al., 2025) or ensemble coding (Park et al., 2011; Nardin et al., 2023). Consistently, our results indicate that studies focusing on single-cell drift inherently depend on the choice of PC detection method. In contrast, population-level analyses offer a more integrative and potentially less biased characterization of hippocampal representational dynamics. In fact, recent findings using such approaches suggest that long-time scale changes in hippocampal representations are not random but preserve underlying population structure over time (Keinath et al., 2022; Sylte et al., 2025).

Population-level analysis makes optimality assumptions about the downstream neural structures reading the population code - for example, decoders are typically trained by minimizing reconstruction errors. Multiple downstream targets may be concerned with distinct aspects of the population code -for example, a readout focusing only on long-term stability may particularly pay attention to SI-PCs, whereas a readout interested in efficient few shot learning may look at all neurons, PCs, and NPCs. The notion that a population may signal distinct information to different targets has recently been put forward under the framework of communication subspaces (Semedo et al., 2019), which are linear subspaces not orthogonal to the weight vector of a linear readout. Thus, distinguishing single cell response properties within a brain area like CA1 may still be important under the assumption that the diversity of information represented in a brain area can be assessed by the diversity of single cell response profiles, and eventually be generated by a diversity of circuit mechanisms. In the case of CA1 place tuning, mechanisms like behavioral time scale plasticity (Bittner et al., 2017), sparse coding (De Almeida et al., 2009), and phase precession (Hussaini et al., 2011) can only be studied if single cell responses are categorized appropriately.

## CRediT authorship contribution statement

**Vladislav Ivantaev**: Data curation, Formal analysis, Investigation, Software, Visualization, Writing – original draft, Writing – review & editing.

**Alireza Chenani**: Conceptualization, Data curation, Investigation, Software.

**Alessio Attardo**: Conceptualization, Data curation, Funding acquisition, Resources, Writing – review & editing.

**Christian Leibold**: Conceptualization, Formal analysis, Funding acquisition, Methodology, Software, Project administration, Supervision, Writing – review & editing.

## Funding

This work was supported by the German Science Association (DFG) under grant numbers AT205/10-1 and LE2250/19-1.

## Declaration of Competing Interest

The authors declare no competing interests.

## Data and code availability

Python 3 code is available from the GITHUB repository https://github.com/ivantaev/pcmethodscomparison. All pre- and fully processed data can be found at the following repository https://gin.g-node.org/ivantaev/Place-cell-detection-data

## Notes

### Competing Interest Statement

The authors have declared no competing interest.

