## Supplementary Information for "Comparison of place field detection methods and their effect on place field stability and drift in mouse dCA1"

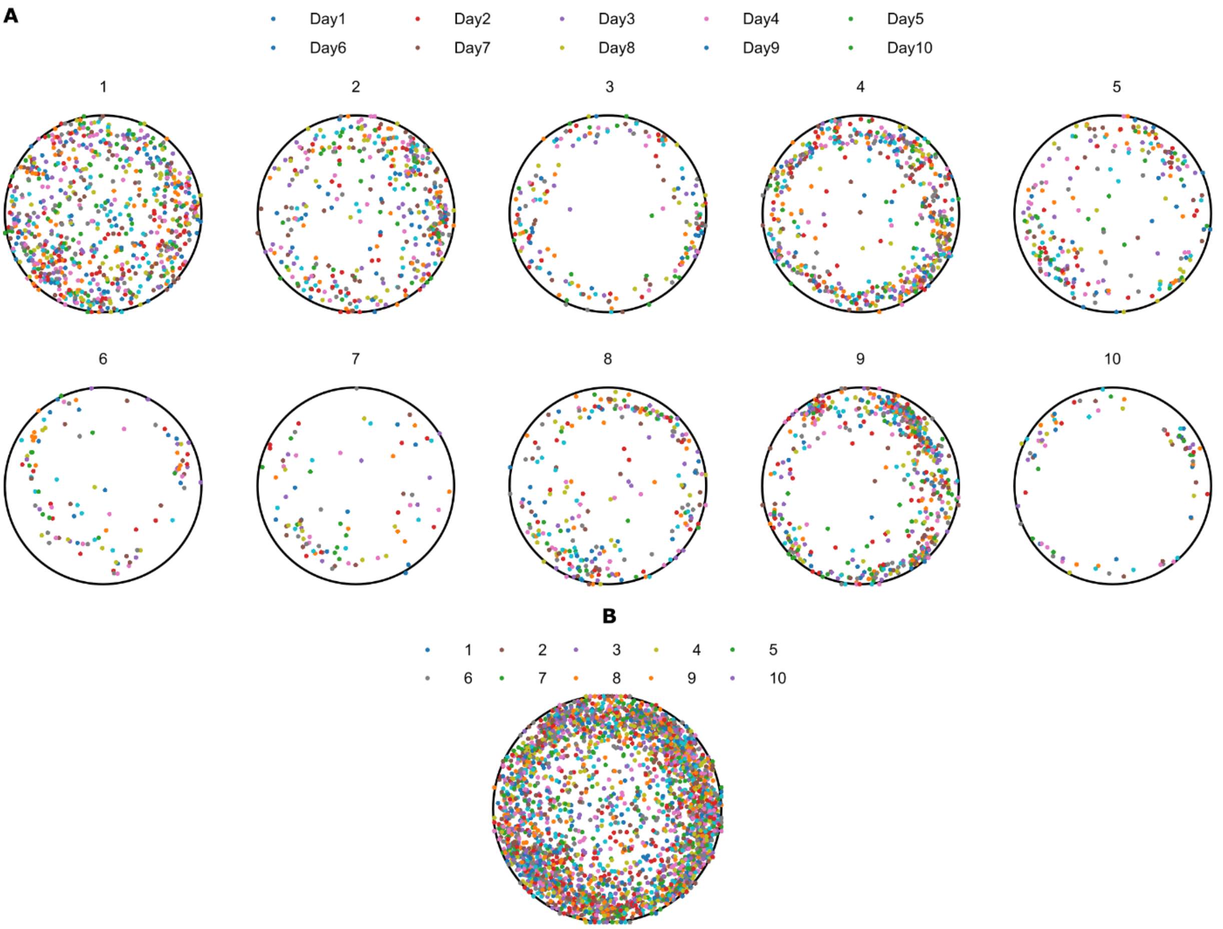


**Supplementary Figure 1. Place cell centroid sampling, consecutive PF CM shifts.**

**A**: CMs of place fields of SI PCs for ten individual mice, each color depicts a different session.

**B**. Same as in **A**, but sampling is plotted for all the animals and color-coded according to an individual mouse.

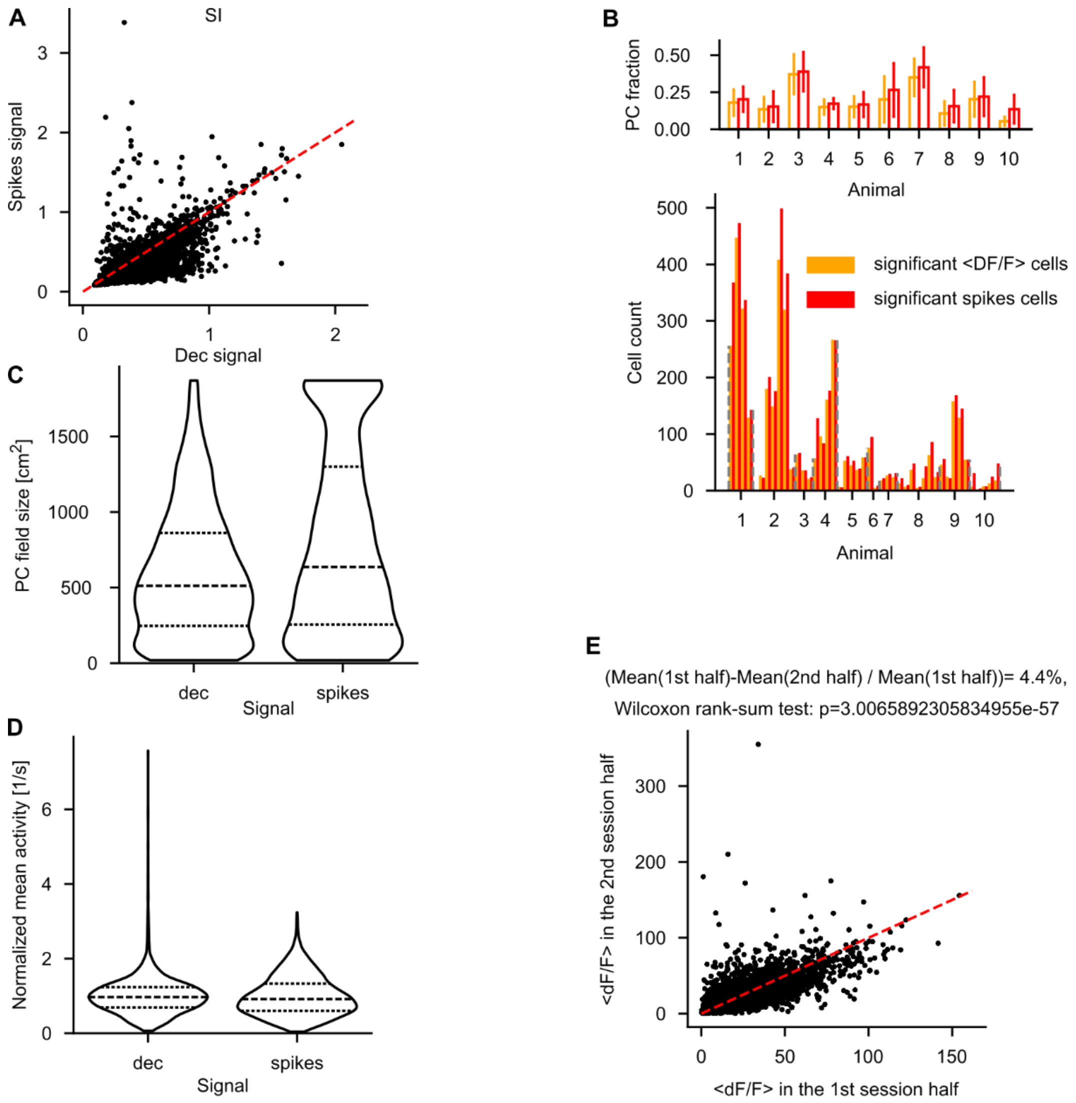


**Supplementary Figure 2. Comparison between Spikes and <DF/F> signal: SI, PC fractions, field sizes.**

**A**: Comparison between the SI content for all cells using either the <DF/F> signal (0.27±0.03 bits) or the spikes signal (0.25±0.15 bits, see **Section 2.2**). Each dot corresponds to a single cell, red dotted line is a unity line.

**B**: Comparison between the PC fraction utilizing the two aforementioned signal types (<DF/F>: 16.6±13.2%, spikes: 20.5±13.7% per session). Each doubled bar corresponds to a single session. Inset: mean count per subject.

**C**: Distributions of placefield sizes of PCs for both signal types (<DF/F>: 590±433 $cm^2$, spikes: 797±622 $cm^2$).

**D**: Distributions of normalized mean PC activities for both signal types (<DF/F>: 1±0.58 1/s, spikes: 1±0.53 1/s). For a fairer comparison, each distribution was normalized by its mean.

**E**: Comparison between the mean fluorescence values in the first and second halves of a session. Each dot corresponds to a single transient, red dotted line is a unity line.

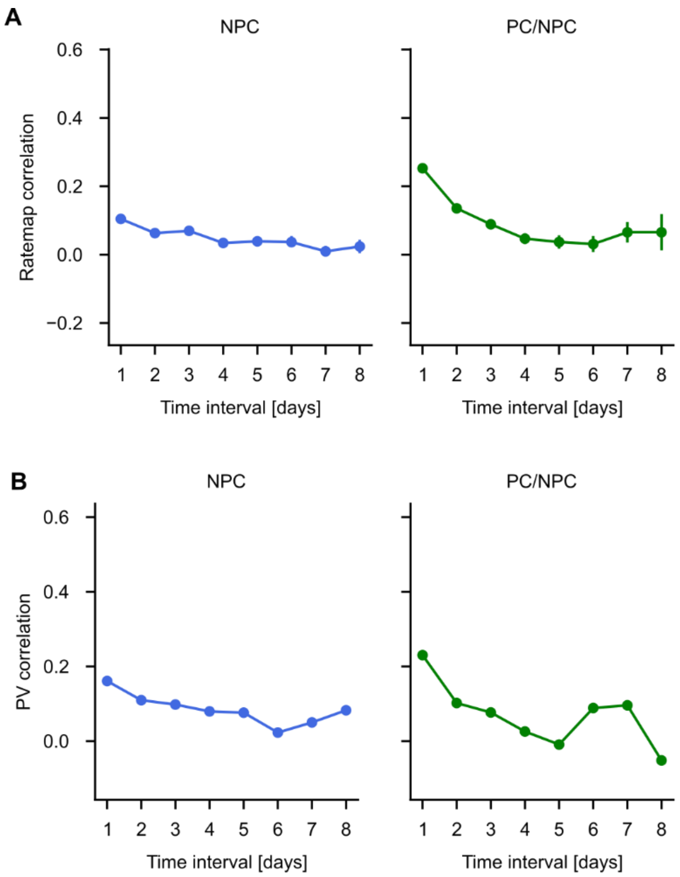


**Supplementary Figure 3. Temporal dynamics of correlations of non-place cells and appearing/disappearing SI-place cells.**

**A**: Average rate map correlations of NPCs and PC/NPCs. Error bars correspond to S.E.M.

**B**: Average population vector correlations of NPCs and PC/NPCs. Error bars correspond to S.E.M.

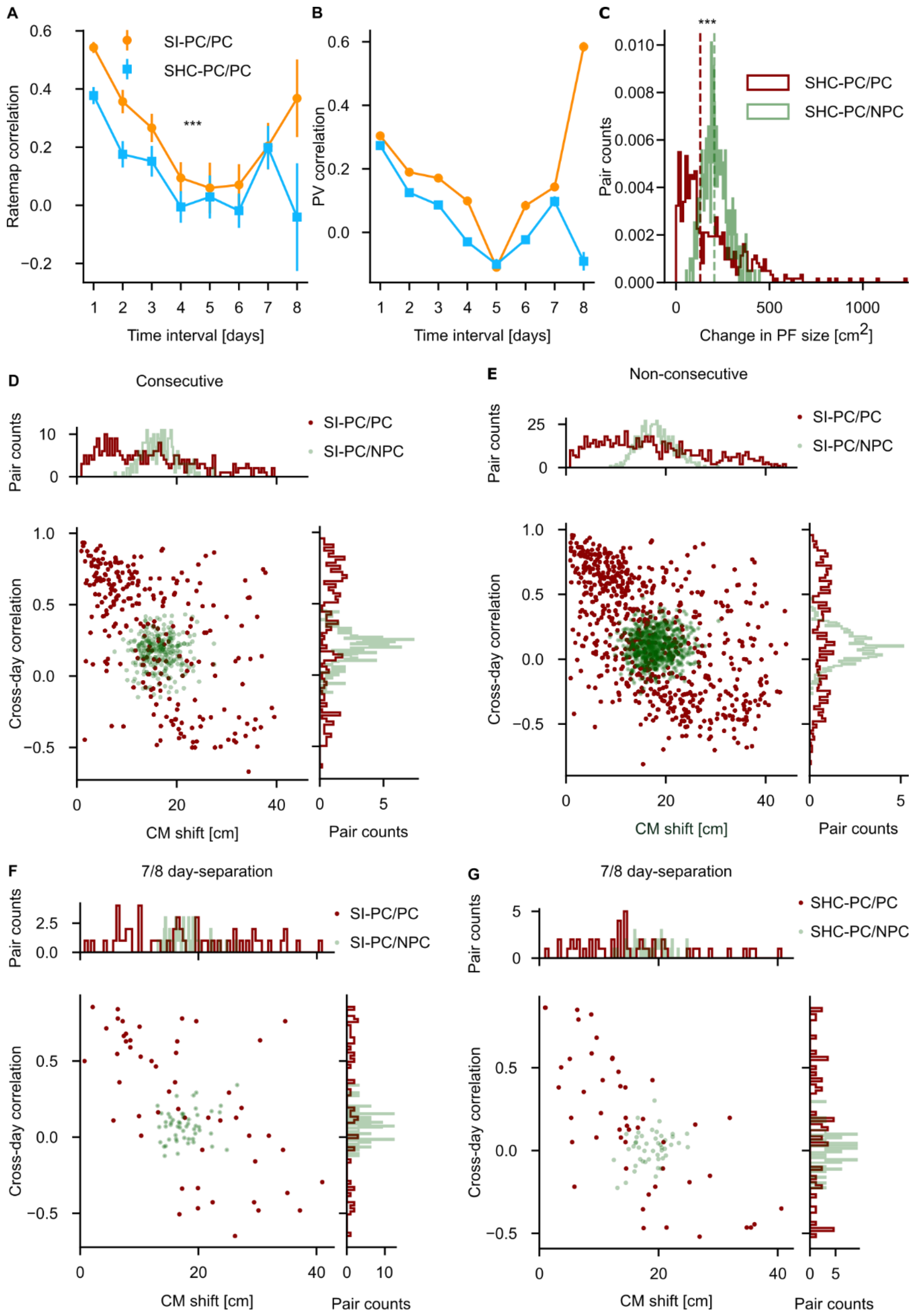


**Supplementary Figure 4. SHC Place cells drift less on average, but change their size more over time than SI place cells.**

**A**: Ratemap correlations as a function of time for PC/PC cell populations detected by SI and SHC methods. Two-way ANOVA (***: $p_{method}$~$10^{-8}$, $F_{method}$=33, $df_{method}$=1; $p_{time}$~$10^{-36}$, $F_{time}$=29, $df_{time}$=7; $p_{intersect}$=0.59, $F_{intersect}$=0.79, $df_{intersect}$=7).

**B**: Same as in **A**, but for the population vector correlation. Two-way ANOVA ($p_{method}$~$10^{-57}$, $F_{method}$=257, $df_{method}$=1; $p_{time}$=0, $F_{time}$=254, $df_{time}$=7; $p_{intersect}$~$10^{-94}$, $F_{intersect}$=66, $df_{intersect}$=7).

**C**: Distribution of changes in PF size for the SHC-PC/PC and down-sampled SHC-PC/NPC populations. Kruskal-Wallis test (***: $p \sim 10^{-15}$, $H=61$, $n_{pc/npc}=502$, $n_{pc/pc}=502$).

**D:** Joint distribution of cross-day rate map correlation vs. CM shift for down-sampled SHC-PC/NPC and SHC-PC/PC populations, tracked for consecutive sessions (ten random down-samples of the SHC-PC/NPC population were performed to match the size of the SHC-PC/PC population). Dots are cell pairs; the dotted line depicts the correlation threshold of 0.3 introduced in **Fig. 2**. Top: marginal distribution of CM shifts; right: marginal distribution of cross-day correlations.

**E:** Same as in **D**, but for the non-consecutive sessions.

**F**: Same as in **D**, but for the session pairs separated by a 7- or 8-day difference.

**G**: Same as in **F**, but for the SHC-PCs.

**A, B**: Orange: SI-PCs; light blue: SHC-PCs. Error bars correspond to S.E.M.

**C, D, E, F, G**: Green: SHC-PC/NPC; red: SHC-PC/PC.

### Supplementary Table 1 | Parameter set used in CaImAn

| Name | Value | Remark |
|---|---|---|
| pw_rigid | False | Performing piecewise-rigid motion correction |
| gSig_filt | (9,9) | Size of high-pass spatial filtering kernel |
| Max_shifts | (25,25) | Maximum allowed rigid shifts |
| Border_nan | ‘copy’ | Treatment of NaN values along the FOV borders |
| Strides | None | |
| overlaps | None | |
| Max_deviation_rigid | 0 | |
| fr | 45 Hz | Imaging frame rate |
| Decay_time | 0.4 s | Decay time of calcium indicator |
| rf | 40 | Half patch size in pixels |
| Stride_cnmf | 20 | Overlap between neighboring patches in pixels |
| gSig | (2,2) | Expected half-size of neurons in pixels |
| gSiz | 9 | Expected size of neurons in pixels (2*gSig+1) |
| Method_init | ‘corr_pnr’ | |
| ssub | 1 | Space down-sampling factor for initialization |
| tsub | 10 | Time down-sampling factor for initialization |
| Ain | None | Possibility to seed with predetermined binary masks |
| gnb | 0 | Number of background components |
| Ring_size_factor | 1.5 | Ring radius |

| | | |
|---|---|---|
| Ssub_B | 2 | Additional background space down-sampling factor |
| Nb_patch | 8 | Number of background components per patch |
| Border_pix | 15 | Assumed FOV border width |
| Min_corr | 0.8 | Minimum value of correlation image for determining candidate components |
| Min_pnr | 5 | Minimum value of psnr image for determining candidate components |
| Merge_thr | 0.7 | Merging threshold of candidate components |
| p | 2 | Order of the autoregressive model |
| Del_duplicates | True | Delete duplicate components in the overlapping regions between neighboring patches |
| Method_deconvolution | 'oasis' | |
| Only_init | True | |
| low_rank_background | None | |
| K | None | |
| Normalize_init | False | |
| Center_psf | True | |
| update_background_components | False | |
| Bas_nonneg | False | |

**Supplementary Table 2 | Sessions considered across subjects**

| Session/subject | 1 | 2 | 3 | 4 | 5 | 6 | 7 | 8 | 9 | 10 |
|---|---|---|---|---|---|---|---|---|---|---|
| 1 | **C/T** | **C/T** | I/A | I/A | T | T | T | **C/T** | **C/T** | R |
| 2 | **C/T** | **C/T** | **C/T** | **C/T** | **C/T** | **C/T** | I/A | I/A | **C/T** | T |
| 3 | R | **C/T** | **C/T** | **C/T** | T | I/A | I/A | I/A | I/A | I/A |
| 4 | **C/T** | **C/T** | T | I/A | **C/T** | I/A | **C/T** | S | S | S |
| 5 | **C/T** | **C/T** | **C/T** | **C/T** | R | T | **C/T** | S | S | S |
| 6 | **C/T** | **C/T** | T | R | R | R | R | S | S | S |
| 7 | R | T | **C/T** | A | **C/T** | **C/T** | R | S | S | S |
| 8 | **C/T** | **C/T** | **C/T** | **C/T** | **C/T** | **C/T** | **C/T** | S | S | S |
| 9 | A | **C/T** | R | **C/T** | **C/T** | **C/T** | **C/T** | I/A | I/A | T |
| 10 | R | I/A | I/A | I/A | T | **C/T** | **C/T** | **C/T** | **C/T** | **C/T** |

Notations. Criteria applied to data from Chenani et al., 2022, in the order of exclusion from the analysis:

A: Incomplete recording (<15 minutes long); not considered.

S: Data from a behavioral condition (see Chenani et al., 2022); not considered.

T: insufficient coverage of the arena (see **Section 2.2**); running trajectory analyzed, but imaging data not considered.

R: bad quality recordings (<60 cells/FOV detected, see **Section 2.2**); CaImAn was applied to a short recording to assess recording quality, but the full session was not considered.

I/A: insufficient footprint alignment between sessions; CaImAn was successfully applied to the full recording that did not match each of the exclusion criteria above. However, due to a change in the FOV and/or ROIs' footprints' quality, the efficiency of the CellReg alignment decreases with the inclusion of the "I/A" session. Hence, all sessions marked as "I/A" were not considered in the final analysis, with the only exception of subjects 3 (Days 6-10) and 10 (Days 2-4) being two subsets of data originally recorded from the same animal (see **Section 2.4**).

C/T: considered, multiday tracked, present in the current analysis.
